# DNA-guided DNA cleavage at moderate temperatures by *Clostridium butyricum* Argonaute

**DOI:** 10.1101/534206

**Authors:** Jorrit W. Hegge, Daan C. Swarts, Stanley D. Chandradoss, Tao Ju Cui, Jeroen Kneppers, Martin Jinek, Chirlmin Joo, John van der Oost

**Affiliations:** Laboratory of Microbiology, Department of Agrotechnology and Food Sciences, Wageningen University, 6708WE Wageningen, The Netherlands; Laboratory of Biochemistry, Department of Agrotechnology and Food Sciences, Wageningen University, 6708WE Wageningen, The Netherlands; Department of Biochemistry, University of Zurich, CH-8057 Zurich, Switzerland.; Kavli Institute of NanoScience, Department of BioNanoScience, Delft University of Technology, 2629HZ Delft, The Netherlands.

## Abstract

Prokaryotic Argonaute proteins (pAgos) constitute a diverse group of endonucleases of which some mediate host defense by utilizing small interfering DNA guides (siDNA) to cleave complementary invading DNA. This activity can be repurposed for programmable DNA cleavage. However, currently characterized DNA-cleaving pAgos require elevated temperatures (≥65°C) for their activity, making them less suitable for applications that require moderate temperatures, such as genome editing. Here we report the functional and structural characterization of the siDNA-guided DNA-targeting pAgo from the mesophilic bacterium *Clostridium butyricum* (*Cb*Ago). CbAgo displays a preference for siDNAs that have a deoxyadenosine at the 5’-end and thymidines in the sub-seed segment (siDNA nucleotides 2-4). Furthermore, *Cb*Ago mediates DNA-guided DNA cleavage of AT-rich double stranded DNA at moderate temperatures (37°C). This study demonstrates that certain pAgos are capable of programmable DNA cleavage at moderate temperatures and thereby expands the scope of the potential pAgo–based applications.

## Introduction

Eukaryotic Argonaute proteins (eAgos) play a key role in RNA interference (RNAi) processes^1–3^. As the core of the multiprotein RNA-induced silencing complex (RISC), eAgos bind small non-coding RNA molecules as guides to direct the RISC complex towards complementary RNA targets^3–5^. Reflecting their physiological function, variation among eAgos is observed with respect to the presence or absence of a catalytic site, and to their potential to interact with other proteins^6^. Depending on the eAgo and on the sequence complementarity between guide and target RNA, eAgo-guide complexes either catalyze endonucleolytic cleavage of the target RNA^7^ or indirectly silence the target RNA by repressing its translation and promoting its degradation through recruitment of additional silencing factors^8^. Independent of the mechanism, eAgo-mediated RNA binding generally results in sequence-specific silencing of gene expression. As such, eAgos can coordinate various cellular processes by regulating intracellular RNA levels.

Prokaryotes also encode Argonaute proteins (pAgos)^9,10^. Various pAgos share a high degree of structural homology with eAgos as both pAgos and eAgos adopt the same four domain (N-PAZ-MID-PIWI) architecture^9–12^. Despite their structural homology, several recently characterized pAgos have distinct functional roles and different guide and/or target preferences compared to eAgos. For example, several pAgos have been implicated in host defense by directly targeting DNA instead of RNA^13–16^. One of the best characterized mechanisms that pAgos utilize is DNA-guided DNA interference, which is demonstrated for pAgos from *Thermus thermophilus* (*Tt*Ago), *Pyrococcus furiosus* (*Pf*Ago), and *Methanocaldococcus jannaschii* (*Mj*Ago)^13–15,17–20^. These pAgos use 5’-end phosphorylated small interfering DNAs (siDNAs) for recognition and successive cleavage of complementary DNA targets. This mechanism enables both *Tt*Ago and *Pf*Ago to mediate host defense against invading nucleic acids. Prokaryotes lack homologs of eukaryotic enzymes that are involved in guide biogenesis^21^. Instead, both *Tt*Ago and *Mj*Ago - besides the canonical siDNA-dependent target cleavage termed ‘slicing’ - exhibit an alternative nuclease activity termed ‘chopping’^14,17^. Chopping facilitates autonomous generation of small DNA fragments from dsDNA substrates. Subsequently, these DNA fragments generated during chopping can serve as siDNAs for canonical slicing^14,17^.

*Tt*Ago and *Pf*Ago can be programmed with short synthetic siDNA which allows them to target and cleave dsDNA sequences of choice *in vitro*^13,15^. This activity has enabled the repurposing of *Pf*Ago as an universal restriction endonuclease for *in vitro* molecular cloning^22^. In addition, a diagnostic *Tt*Ago-based application termed NAVIGATER (Nucleic Acid enrichment Via DNA Guided Argonaute from *Thermus thermophilus*) was developed which enables enhanced detection of rare nucleic acids with single nucleotide precision^23^. In analogy with the now commonly used CRISPR-Cas9 and CRISPR-Cas12a enzymes^24–26^, it has also been suggested that pAgos could be repurposed as next-generation genome editing tools^27^. However, due to the thermophilic nature (optimum activity temperature ≥65°C) and low levels of endonuclease activity at the relevant temperatures (20-37°C), it is unlikely that the well-studied *Tt*Ago, *Pf*Ago and *Mj*Ago are suitable for genome editing. The quest for a pAgo that can cleave dsDNA at moderate temperatures has resulted in the characterization of the Argonaute protein from *Natronobacterium gregory* (*Ng*Ago), which was claimed to be the first pAgo suitable for genome editing purposes^28^. However, the study reporting this application has been retracted after a series of reproducibility issues^28–30^. Instead, it has been suggested that *Ng*Ago targets RNA rather than DNA^31^.

Although considerable efforts have been made to elucidate the mechanisms and biological roles of pAgos, efforts have mainly focused on pAgo variants from (hyper)thermophiles. This has left a large group of mesophilic pAgos unexplored. We here report the characterization of the Argonaute protein from the mesophilic bacterium *Clostridium butyricum* (*Cb*Ago). We demonstrate that *Cb*Ago can utilize siDNA guides to cleave both ssDNA and dsDNA targets at moderate temperatures (37°C). In addition, we have elucidated the macromolecular structure of *Cb*Ago in complex with a siDNA guide and complementary ssDNA target in a catalytically competent state. *Cb*Ago displays an unusual preference for siDNAs with a deoxyadenosine at the 5’-end and thymidines in the sub-seed segment (siDNA nt 2-4). The programmable DNA endonuclease activity of *Cb*Ago provides a foundation for the development of pAgo-based applications at moderate temperatures..

## Materials and methods

### Plasmid construction

The *Cb*Ago gene was codon harmonized for E.coli Bl21 (DE3) and inserted into a pET-His6 MBP TEV cloning vector (obtained from the UC Berkeley MacroLab, Addgene #29656) using ligation independent cloning (LIC) using oligonucleotides oDS067 and oDS068 (Table S1) to generate a protein expression construct that encodes the *Cb*Ago polypeptide sequence fused to an N-terminal tag comprising a hexahistidine sequence, a maltose binding protein (MBP) and a Tobacco Etch Virus (TEV) protease cleavage site.

### Generation of the Double mutant

*Cb*Ago double mutant (D541A, D611A) was generated using an adapted Quick Directed Mutagenesis Kit instruction manual (Stratagene). The primers were designed using the web-based program primerX (http://bioinformatics.org/primerx).

### *Cb*Ago expression and purification

The *Cb*Ago WT and DM proteins were expressed in E.coli Bl21(DE3) Rosetta™ 2 (Novagen). Cultures were grown at 37°C in LB medium containing 50 μg ml-1 kanamycin and 34 μg ml-1 chloramphenicol until an OD600nm of 0.7 was reached. *Cb*Ago expression was induced by addition of isopropyl β-D-1-thiogalactopyranoside (IPTG) to a final concentration of 0.1 mM. During the expression cells were incubated at 18°C for 16 hours with continues shaking. Cells were harvested by centrifugation and lysed by sonication (Bandelin, Sonopuls. 30% power, 1s on/2s off for 5min) in lysis buffer containing 20 mM Tris-HCl pH 7.5, 250 mM NaCl, 5 mM imidazole, supplemented with a EDTA free protease inhibitor cocktail tablet (Roche). The soluble fraction of the lysate was loaded on a nickel column (HisTrap Hp, GE healthcare). The column was extensively washed with wash buffer containing 20 mM Tris-HCl pH 7.5, 250 mM NaCl and 30 mM imidazole. Bound protein was eluted by increasing the concentration of imidazole in the wash buffer to 250 mM. The eluted protein was dialysed at 4°C overnight against 20 mM HEPES pH 7.5, 250 mM KCl, and 1mM dithiothreitol (DTT) in the presence of 1mg TEV protease (expressed and purified according to Tropea et al. 2009^55^) to cleave of the His6-MBP tag. Next the cleaved protein was diluted in 20mM HEPES pH 7.5 to lower the final salt concentration to 125 mM KCl. The diluted protein was applied to a heparin column (HiTrap Heparin HP, GE Healthcare), washed with 20 mM HEPES pH 7.5, 125 mM KCl and eluted with a linear gradient of 0.125-2 M KCl. Next, the eluted protein was loaded onto a size exclusion column (Superdex 200 16/600 column, GE Healthcare) and eluted with 20 mM HEPES pH 7.5, 500mM KCl and 1 mM DTT. Purified *Cb*Ago protein was diluted in size exclusion buffer to a final concentration of 5 μM. Aliquots were flash frozen in liquid nitrogen and stored at −80°C.

### Co-purification nucleic acids

To 500 pmoles of purified *Cb*Ago in SEC buffer CaCl2 and proteinase K (Ambion) were added to final concentrations of 5 mM CaCl2 and 250 μg/mL proteinase K. The sample was incubated for 4 hours at 65°C. The nucleic acids were separated from the organic fraction by adding Roti phenol/chloroform/isoamyl alcohol pH 7.5-8.0 in a 1:1 ratio. The top layer was isolated and nucleic acids were precipitated using ethanol precipitation by adding 99% ethanol in a 1:2 ratio supplied with 0.5% Linear polymerized acrylamide as a carrier. This mixture was incubated overnight at −20°C and centrifuged in a table centrifuge at 16,000 g for 30 min. Next, the nucleic acids pellet was washed with 70% ethanol and solved in 50 μL MilliQ water. The purified nucleic acids were treated with either 100 μg/mL RNase A (Thermo), 2 units DNase I (NEB) or both for 1 hour at 37°C and resolved on a denaturing urea polyacrylamide gel (15%) and stained with SYBR gold.

### Single stranded Activity assays

Unless stated otherwise 5 pmoles of each *Cb*Ago, siDNA and target were mixed in a ratio of 1:1:1, in 2x reaction buffer containing 20 mM Tris-HCl (pH 7.5) supplemented with 500 μM MnCl^2+^. The target was added after the *Cb*Ago and siDNA had been incubation for 15 min at 37°C. Then the complete reaction mixture was incubated for 1 hour at 37°C. The reaction was terminated by adding 2x RNA loading dye (95% Formamide, 0.025% bromophenol blue, 5 mM ETDA) and heating it for 5 minutes at 95°C. After this the samples were resolved on a 20% denaturing (7 M Urea) polyacrylamide gel. The gel was stained with SYBR gold nucleic acid stain (Invitrogen) and imaged using a G:BOX Chemi imager (Syngene).

### Double stranded Activity assay

In two half reactions 12.5 pmoles of *Cb*Ago was loaded with either 12.5pmoles of forward or reverse siDNA in reaction buffer containing 10 mM Tris-HCl, 10 μg/ml BSA, 250 μM MnCl2. The half reactions were incubated for 15 min at 37°C. Next, both half reactions were mixed together and 120 ng target plasmid was added after which the mixture was incubated for 1 hour of 37°C. After the incubation the target plasmid was purified from the mixture using a DNA clean and concentrate kit (DNA Clean & Concentrator™-5, Zymogen) via the supplied protocol. The purified plasmid was subsequently cut using either EcoRI-HF (NEB) or SapI-HF (NEB) in Cutsmart buffer (NEB) for 30 min at 37°C. A 6x DNA loading dye (NEB) was added to the plasmid sample prior to resolving it on a 0.7% agarose gel stained with SYBR gold (Invitrogen).

### Crystallization

To reconstitute the *Cb*Ago DM-siDNA-target DNA complex, siDNA and target DNA were pre-mixed at a 1:1 ratio, heated to 95°C, and slowly cooled to room temperature. The formed dsDNA duplex (0.5M) was mixed with *Cb*Ago DM in SEC buffer at a 1:1:4 ratio (*Cb*Ago DM:duplex DNA), and MgCl2 was added to a final concentration of 5 mM. The sample was incubated for 15 minutes at 20°C to allow complex formation. The complex was crystallized at 20°C using the hanging drop vapour diffusion method by mixing equal volumes of complex and reservoir solution. Initial crystals were obtained at a *Cb*Ago DM concentration of 5 mg/ml with a reservoir solution consisting of 4 M Sodium Formate. Data was collected from crystals grown obtained using a complex concentration of 4.3 mg/ml and reservoir solution containing 3.8 M Sodium Formate and 5 mM NiCl2 at 20°C. For cryoprotection, crystals were transferred to a drop of reservoir solution and flash-cooled in liquid nitrogen.

X-ray diffraction data were measured at beamline X06DA (PXIII) of the Swiss Light Source (Paul Scherrer Institute, Villigen, Switzerland). Data were indexed, integrated, and scaled using AutoPROC (Vonrhein et al (2011)). Crystals of the *Cb*Ago-siDNA-target DNA complex diffracted to a resolution of 3.55 Å and belonged to space group *P*63 2 2, with one copy of the complex in the asymmetric unit. The structure was solved by molecular replacement using Phaser-MR (McCoy et al., 2007). As search model, the structure of *Tt*Ago in complex with guide and target DNA strands (PDB: 5GQ9) was used after removing loops and truncating amino acid side chains. Phases obtained using the initial molecular replacement solution were improved by density modification using phenix.resolve (Terwilliger, 2004) and phenix.morph_model (Terwilliger et al., 2013). The atomic model was built manually in Coot (Emsley et al., 2010) and refined using phenix.refine (Afonine et al., 2012). The final binary complex model contains *Cb*Ago residues 1-463 and 466-748, guide DNA residues 1–16, and target DNA residues (−18)–(−1).

### Structure analysis

Core Root Means Square Deviations (rmsd) of structure alignments were calculated using Coot SSM superpose (Krissinel et al 2004). Intramolecular interactions were analysed using PDBePISA (Krissinel and Henrick, 2007). Figures were generated using PyMOL (Schrödinger).

### Single-Molecule Experimental Set-Up

Single-molecule fluorescence FRET measurements were performed with a prism-type total internal reflection fluorescence microscope. Cy3 and Cy5 molecules were excited with 532 nm and 637 nm wavelength, respectively. Resulting Cy3 and Cy5 fluorescence signal was collected through a 60X water immersion objective (UplanSApo, Olympus) with an inverted microscope (IX73, Olympus) and split by a dichroic mirror (635dcxr, Chroma). Scattered laser light was blocked out by a triple notch filter (NF01-488/532/635, Semrock). The Cy3 and Cy5 signals were recorded using a EM-CCD camera (iXon Ultra, DU-897U-CS0-#BV, Andor Technology) with exposure time 0.1 s. All single-molecule experiments were done at room temperature (22 ± 2C).

### Fluorescent DNA and RNA preparation

The RNAs with amine-modification (amino-modifier C6-U phosphoramidite, 10-3039, Glen Research) were purchased from STPharm (South Korea) and DNAs with amine-modification (internal amino modifier iAmMC6T) Ella biotech (Germany). The guide and target strands were labeled with donor (Cy3) and acceptor (Cy5), respectively, using the NHS-ester form of Cy dyes (GE Healthcare). 2012).1 μL of 1 mM of DNA/RNA dissolved in MilliQ H20 is added to 5 μL labeling buffer of (freshly prepared) sodiumtetraborate (380 mg/10mL, pH 8.5). 1 μL of 20 mM dye (1 mg in 56 μL DMSO) is added and incubated overnight at room temperature in the dark, followed by washing and ethanol precipitation. The labeling efficiency was ~100%.

### Single-molecule sample preparation

A microfluidic chamber was incubated with 20 μL Streptavidin (0.1 mg/mL, Sigma) for 30 sec. Unbound Streptavidin was washed with 100 μL of buffer T50 (10 mM Tris-HCl [pH8.0], 50 mM NaCl buffer). The fifty microliters of 50 pM acceptor-labelled target construct were introduced into the chamber and incubated for 1 min. Unbound labeled constructs were washed with 100 μL of buffer T50. The CbAgo binary complex was formed by incubating 10 nM purified CbAgo with 1 nM of donor-labeled guide in a buffer containing 50 mM Tris-HCl [pH 8.0] (Ambion), 1mM MnCl2, and 100 mM NaCl (Ambion) at 37°C for 20 min. For binding rate (kon) measurements, the binary complex was introduced into the fluidics chamber using syringe during the measurement. The experiments were performed at the room temperature (23 ± 1°C).

For fluorescence Guide Loading Experiments before immobilizing CbAgo on the single-molecule surface, 1 μL of 5 μM His-tagged apo-CbAgo was incubated with 1 μL of 1 μg/ml biotinylated anti-6x His antibody (Abcam) for 10 min. Afterward, the mixture was diluted 500x in T50 and 50 μL were loaded in the microfluidic channel for 30 s incubation, followed by washing with 100 μL of T50 buffer. Cy3-labeled ssDNA (0.1) was applied to the microfluidic chamber in imaging buffer (50 mM Tris-HCl pH 8.0, 100 mM NaCl, 1 mM MnCl2, 1 mM Trolox ((±)-6-Hydroxy-2,5,7,8-tetramethylchromane-2-carboxylic acid), supplemented with an oxygen-scavenging system (0.5 mg/mL glucose oxidase (Sigma), 85 mg/mL catalase (Merck), and 0.8% (v/v) glucose (Sigma)).

### Single-molecule data acquisition and analysis

CCD images of time resolution 0.1 or 0.3 sec were recorded, and time traces were extracted from the CCD image series using IDL (ITT Visual Information Solution). Co-localization between Cy3 and Cy5 signals was carried out with a custom-made mapping algorithm written in IDL. The extracted time traces were processed using Matlab (MathWorks) and Origin (Origin Lab).

The binding rate (*k*_*on*_) was determined by first measuring the time between when CbAgo binary complex was introduced to a microfluidic chamber and when the first CbAgo-guide docked to a target; and then fitting the time distribution with a single-exponential growth curve, 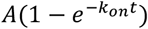. The dissociation rate was estimated by measuring the dwell time of a binding event. A dwell time distribution was fitted by single-exponential decay curve (*Ae*^−*t*/Δ*τ*^).

### Fluorescence competition experiments

MBP-tagged CbAgo was immobilized on the quartz surface using an anti-MBP antibody. An equimolar mixture of let7 DNA guide (Cy3 labeled) and let7 RNA guide (Cy5 labeled) in imaging buffer was introduced to the microfluidic chamber. After 5 minutes, 10 snapshots of independent fields of view with simultaneous illumination were collected to estimate the amount of guide molecules bound to protein. Movies were taken for 200 s (2000 frames) at continuous illumination of Cy3 and Cy5 molecules to determine the dwell times of the binding events. Dwell times were binned in a histogram and fitted with a single exponential decay curve.

### FRET targeting experiments of ATTT and AAAA guide target combinations

100 pM of target construct annealed with biotin handle were flushed in the microfluidic chamber. After incubation of 1 min, the microfluidic chamber was rinsed with 100 μL T50 buffer. 10 nM of apo-CbAgo was loaded with 1 nM of ATTT seed DNA guide or with AAAA seed DNA guide at 37°C for 30 minutes in imaging buffer after which the mixture is introduced inside the microfluidic chamber. Movies of 200 s were taken at continuous illumination of the Cy3 signal. Site specific protein target interactions were identified as FRET signals and were further analysed.

## Results

### *Cb*Ago mediates siDNA-guided ssDNA cleavage

*Cb*Ago was successfully expressed in *E. coli* from a codon-optimized gene using a T7-based pET expression system and purified (Supplementary Figure S1A). To determine the guide and target binding characteristics of *Cb*Ago, we performed single-molecule experiments using Förster resonance energy transfer (FRET). We immobilized either Cy5-labeled single stranded RNA or DNA targets (FRET acceptor) on a polymer-coated quartz surface (Figure 1A). Next we introduced *Cb*Ago in complex with either a Cy3-labeled siRNA or siDNA guide (FRET donor) and recorded the interactions. Strikingly, *Cb*Ago could utilize both siRNAs and siDNAs to bind DNA or RNA targets (Figure 1B). To test which guide is preferentially bound by *Cb*Ago we performed a competition assay in which *Cb*Ago was immobilized into the microfluidic chamber, and an equimolar mixture of siDNA and siRNAs was introduced. While only short-lived interactions (average dwell time: 0.48 seconds) were observed for siRNA, siDNA was strongly bound (average dwell time: 44 seconds) by *Cb*Ago (Figure 1C). This results suggests that *Cb*Ago utilizes siDNA rather than siRNA as a guide.

**Figure 1.**
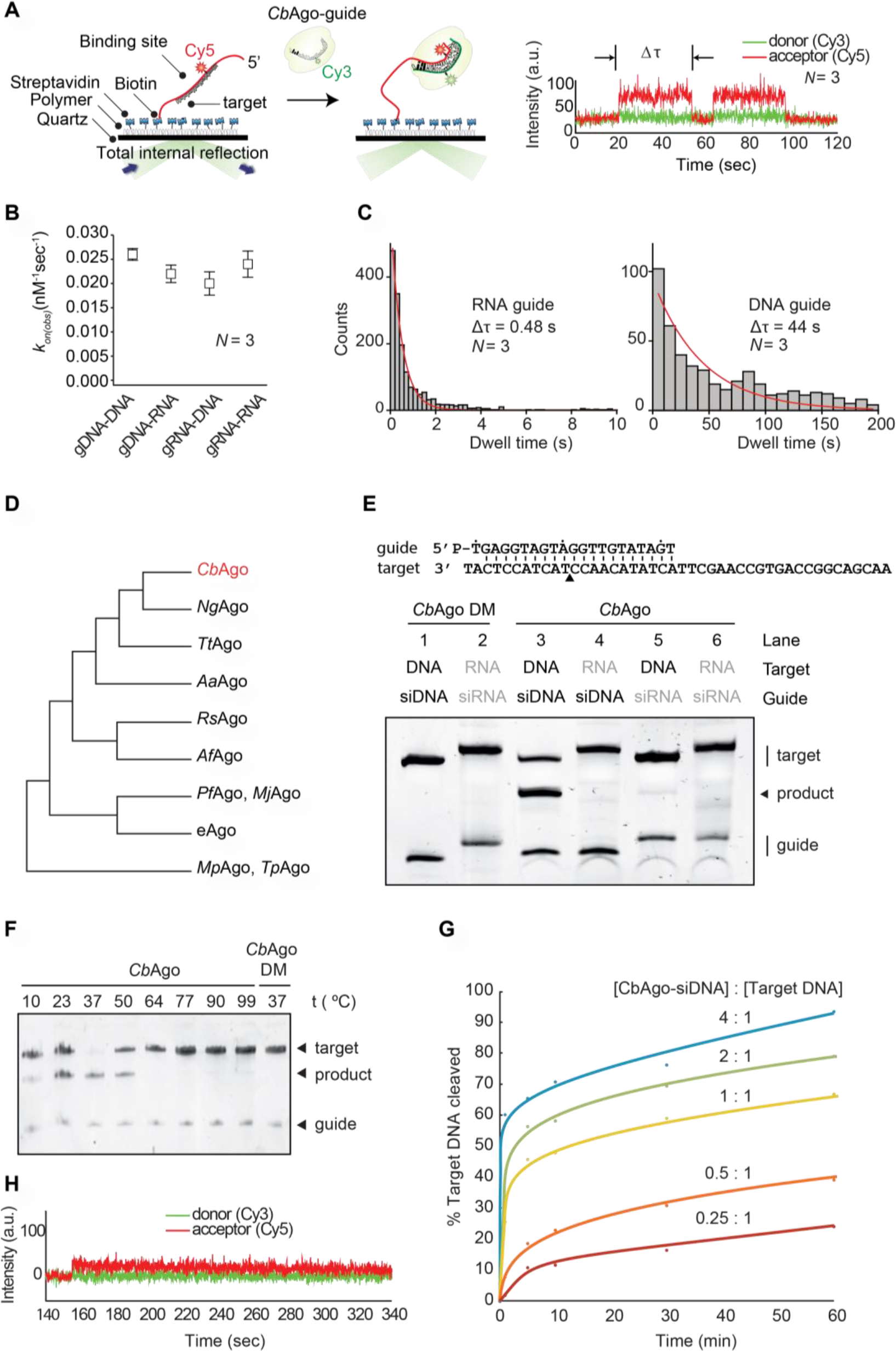
*Cb*Ago exhibits DNA-guided DNA endonuclease activity at 37°C. (**A**) Left: Overview of the single molecule assay to determine the binding characteristics of *Cb*Ago. Right: FRET diagram of a *Cb*Ago-siDNA complex that has 3 complementary base pairs (2-4nt) to the DNA target. Indicated is the dwell time (Δτ). (**B**) Comparison of the binding rates (k_on_) of *Cb*Ago in complex with siDNA or siRNA to bind DNA or RNA targets. The rates are similar for each nucleic acid type guide and target. *N* is the number of base paired nucleotides. (**C**) Dwell time histograms showing CbAgo preferentially binds siDNAs in siDNA-siRNA competition experiments. (**D**) Schematic phylogenetic tree of characterized pAgos. (**E**) *Cb*Ago exhibits DNA-guided DNA endonuclease activity. Upper panel: Sequence of the synthetic let7 miRNA-based siDNA guide and target DNA sequences that were used for the *in vitro* assays. Lower panel: *Cb*Ago, guides and targets were mixed in a 1:1:1 molar ratio and incubated for 1 h at 37°C. Catalytic mutant CbAgoDM was used as a control. Cleavage products were analysed by denaturing polyacrylamide electrophoresis. (**F**) *Cb*Ago displays highest activity at 37°C. *Cb*Ago and siDNA were mixed and pre-incubated at various temperatures for 10 minutes. Next, target DNA was added and the sample was incubated for 1 h at the same temperature. *Cb*AgoDM was used as a control. Cleavage products were analysed by denaturing polyacrylamide electrophoresis. (**G**) Quantified data of a *Cb*Ago-mediated siDNA-guided ssDNA cleavage turnover experiment using 5 pmol target DNA and increasing concentrations of *Cb*Ago-siDNA (1.25-20 pmol). (**H**) FRET diagram showing that a cleavage compatible *Cb*Ago-siDNA remains bound to a fully complementary target DNA (*N*=21) during the entire the measurement (340 seconds).

*Cb*Ago is phylogenetically closest related to the clade of halobacterial pAgos, among which also pAgo from *Natronobacterium gregoryi* (*Ng*Ago) can be found (Figure 1D and Supplementary Figure S2). A multiple sequence alignment of *Cb*Ago with other pAgos (Supplementary Figure S1B) suggests that *Cb*Ago contains the conserved DEDX catalytic residues (where X can be a D, H or N) which are essential for nuclease activity in ‘slicing’ Agos^32^. In the case of *Cb*Ago, this concerns residues D541, E577, D611 and D727.

To confirm whether *Cb*Ago indeed is an active nuclease, we performed *in vitro* activity assays in which *Cb*Ago was loaded with either synthetic siDNAs or siRNAs (21 nucleotides in length). Next the complexes were incubated at 37°C with 45-nucleotide complementary single stranded RNA or DNA target oligonucleotides. While no activity was found in any of the combinations in which siRNAs or target RNAs were used, *Cb*Ago was able to cleave target DNAs in a siDNA-dependent manner (Figure 1E). In agreement with the predicted DEDD catalytic site (Supplementary Figure S1B), alanine substitutions of two of aspartic acids (D541A, D611A) in the expected catalytic tetrad abolished the nuclease activity, demonstrating that the observed siDNA-guided ssDNA endonucleolytic activity was indeed catalyzed by the DEDD catalytic site. To further investigate the full temperature range at which *Cb*Ago is active, we performed additional cleavage assays at temperatures ranging from 10-95°C. While *Cb*Ago displayed the highest activity at its physiologically relevant temperature (37°C), *Cb*Ago also catalyzed siDNA-guided target DNA cleavage at temperatures as low as 10°C and as high as 50°C (Figure 1F).

When *Cb*Ago-siDNA complexes and target ssDNA substrates (45nt) were mixed in equimolar amounts, cleavage of the target DNA was not complete after one hour incubation (Figure 1E). Therefore, we investigated the substrate turnover kinetics of *Cb*Ago by monitoring the cleavage assays in a time course using variable *Cb*Ago:siDNA:target DNA ratios (Figure 1G). A rapid burst of activity was observed during the first minute, likely indicating the first target binding and cleavage event. This stage was followed by a slow steady state, suggesting that under these conditions the *Cb*Ago-siDNA complex slowly dissociates from the cleaved target DNA product before being able to bind and cleave a new target DNA strand. The cleavage kinetics were confirmed using single-molecule assays which demonstrated that the *Cb*Ago-siDNA complex remains bound to the DNA target (*N*=21) for several minutes (Figure 1H), which prevents *Cb*Ago-siDNA complexes from binding and cleaving new DNA targets. Thus, while CbAgo functions as a multi-turnover nuclease enzyme, its steady-state rate is limited by product release.

### Structure of *Cb*Ago in the cleavage-competent conformation

To investigate the molecular architecture of *Cb*Ago in light of its biochemical activity, we crystallized *Cb*AgoDM in complex with both a 21-nt siDNA and a 19-nt DNA target, and solved the structure of the complex at 3.54 Å resolution (Figure 2 and Table S1). Like other Agos, *Cb*Ago adopts a bilobed conformation in which one of the lobes comprises the N-terminal, linker L1, and PAZ domains, which are linked by linker L2 to the other lobe comprising the MID and PIWI domains. Nucleotides 2-16 of the siDNA constitute a 15 base-pairs A-form-like duplex with the target DNA (Figure 2A). The 5’-terminal nucleotide of the siDNA is anchored in the MID domain pocket, where the 5’-phosphate group of the siDNA makes numerous interactions with MID domain residues and the C-terminal carboxyl group of CbAgo (Supplementary Figure S3). To test whether the interactions with the 5’-phosphate group of the siDNA are important for *Cb*Ago activity, we performed target DNA cleavage assays in which we used siDNAs with a 5’ phosphate or a 5’ hydroxyl group (Supplementary Figure S4). As observed for other pAgos^33,34^, *Cb*Ago is able to utilize both siDNAs for target DNA cleavage, but it cleaves target DNA much more efficiently when the siDNA contains a 5’-phosphate group. This is in agreement with the siDNA-protein interactions observed in the crystal structure. Furthermore, the backbone phosphates of the siDNA seed segment form hydrogen-bonding and ionic interactions with specific residues in the MID, PIWI and L1 domains (Supplementary Figure S3). At the distal end of the siDNA-target DNA duplex, the N-domain residue His35 caps the duplex by stacking onto the last base pair. After this point, the remaining 3’-terminal nucleotides of the siDNA are unordered, while the target DNA bends away from the duplex and enters the cleft between the N-terminal and PAZ domains. In agreement with other ternary pAgo complexes^18,35,36^, the PAZ domain pocket, which normally binds the 3’ end of the guide in a binary Ago-guide complex, is empty.

**Figure 2.**
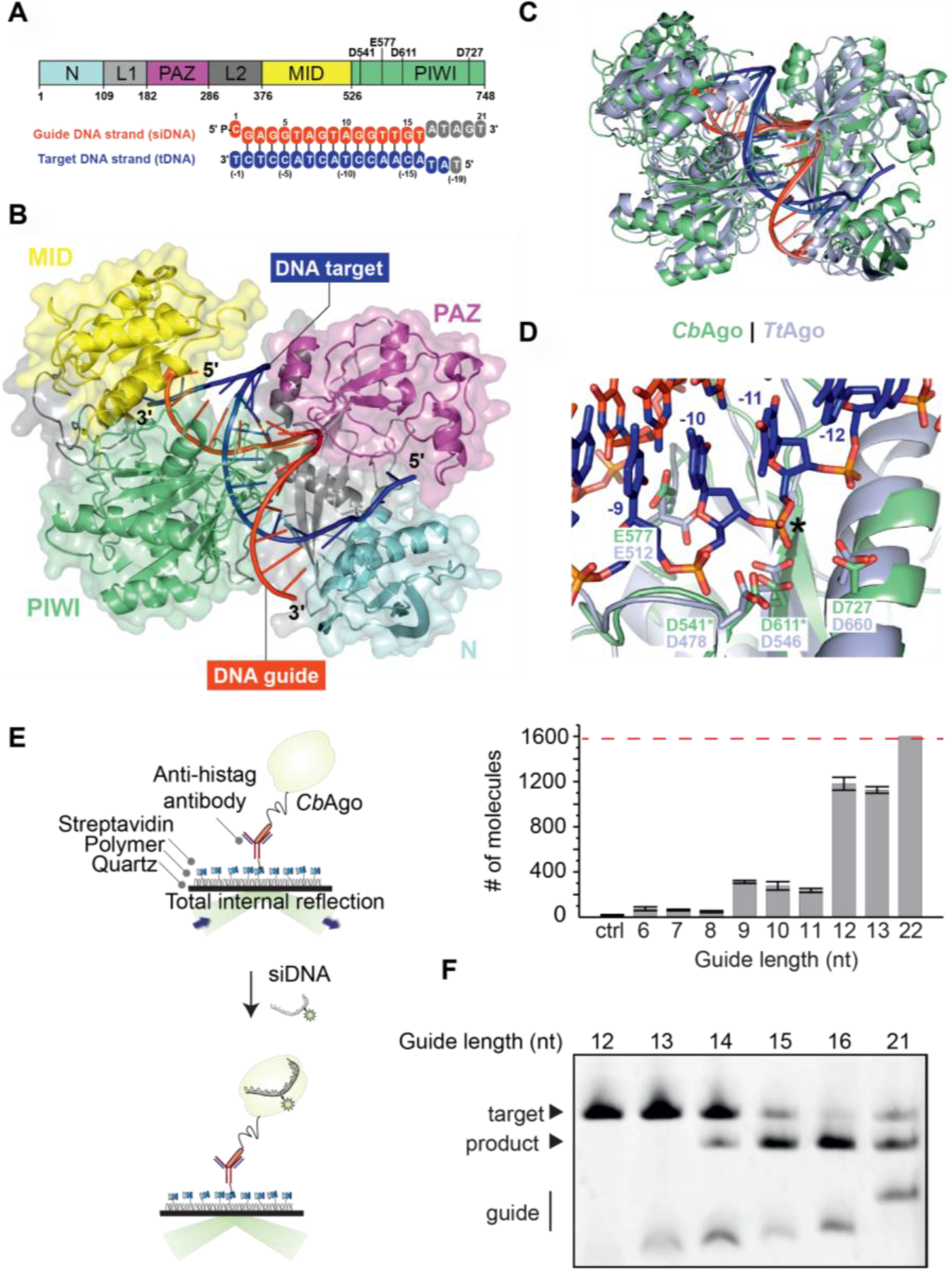
Structure of *Cb*Ago in complex with a siDNA and a DNA target. (**A**) Upper panel: Schematic diagram of the domain organization of *Cb*Ago. L1 and L2 are linker domains. Lower panel: Sequences of the siDNA (red) and target DNA (blue). Nucleotides that are unordered in the structure are coloured grey. See also Table S1. (**B**) Overall structure of the *Cb*Ago-siDNA-target DNA complex. Domains are coloured according to the colour scheme in panel A. (**C**) Structural alignment of *Cb*Ago (green) and *Tt*Ago (light purple; PDB: 4NCB). Core Root Mean Square Deviation of 3.0 Å over 563 residues. (**D**) Close-up view of the aligned DDED catalytic sites of *Cb*Ago (green) and *Tt*Ago (light purple; PDB: 4NCB). Modelled side chains of D541 and D611 in *Cb*Ago are indicated with green asterisks. The glutamate finger of both pAgos (E512 in *Tt*Ago or E577 in *Cb*Ago) are inserted into the catalytic site. The scissile phosphate between nucleotide −10 and −11 of the target DNA strand (blue) is indicated with a black asterisk. (**E**) Total internal reflection microscopy (TIRM) was used to determine the minimal length for siDNA to be bound by *Cb*Ago. Left panel: Graphical overview of the TIRM method. Right panel: Histogram with TIRM results demonstrated that synthetic siDNAs of at least 12nt in length are efficiently bound by *Cb*Ago. The red line indicates the total number of countable molecules within the microscope image. The raw microscope images are given in Supplementary Figure S5. (**F**) *Cb*Ago mediates target DNA cleavage with siDNAs as short as 14 nucleotides. *Cb*Ago was incubated with siDNA and target DNA in a 1:1:1 ratio. Cleavage products were analysed by denaturing polyacrylamide electrophoresis.

*Cb*Ago is phylogenetically closely related to *Tt*Ago (Figure 1D). However, *Cb*Ago is 63 amino acids (9.2%) longer than *Tt*Ago (748 amino acids vs. 685 amino acids) and *Cb*Ago and *Tt*Ago share only 23% sequence identity. Superposition of the *Cb*Ago complex structure with the structure of *Tt*Ago bound to a siDNA and DNA target (PDB: 4NCB) (Figure 2C) reveals that the macromolecular architecture and conformation of these *Tt*Ago and *Cb*Ago structures are highly similar (Core root mean square deviation of 3.0 Å over 563 residues), with differences found mostly in the loop regions. This agrees with the fact that loops of thermostable proteins are generally more compact and shorter^37,38^. In the *Tt*Ago structure, which is thought to represent a catalytically competent state, a ‘glutamate finger’ side chain (Glu512^*Tt*Ago^) is inserted into the catalytic site completing the catalytic DDED tetrad^35^. Similarly, the corresponding residue in *Cb*Ago (Glu577) is located within a flexible loop and is positioned near the other catalytic residues (Figure 2D; Asp541, Asp611, and Asp727). All pAgos and eAgos characterized to date cleave the target strand in between nucleotide 10 and 11 of the target strand. In line with the consensus, the catalytic residues of CbAgo perfectly align with the scissile phosphate linking these nucleotides in our structure (Figure 2D). This observation implies that this structure represents the cleavage competent conformation of *Cb*Ago.

Only 15 siDNA-target DNA base pairs are formed in the complex, which suggests that additional siDNA-target DNA binding is not essential for target DNA cleavage. To determine the minimum siDNA length that *Cb*Ago requires for target binding, we performed single-molecule fluorescence assays. First, *Cb*Ago was immobilized on a surface and next it was incubated with 5’-phosphorylated Cy3-labelled siDNAs (Figure 2E). These assays demonstrate that *Cb*Ago can bind siDNAs with a minimal length of 12 nucleotides. Next, we determined the minimum siDNA length for CbAgo-siDNA mediated target DNA cleavage (Figure 2F). In line with the observation that the *Cb*Ago adopts a cleavage-competent confirmation when only 14 base pairs are formed, *Cb*Ago can cleave target DNAs when programmed with siDNAs as short as 14 nt (forming 13 siDNA-target DNA base pairs) under the tested conditions. This resembles the activity of *Pf*Ago, *Mj*Ago, and *Mp*Ago, which require siDNAs with a minimal length of 15 nt to catalyze target DNA cleavage^14,15,34^. Only *Tt*Ago has been reported to mediate target DNA cleavage with siDNAs as short as 9 nt^12^.

### CbAgo associates with plasmid-derived siDNAs in vivo

It has previously been demonstrated that certain pAgos co-purify with their guides and/or targets during heterologous expression in *Escherichia coli*^13,16^. To determine whether *Cb*Ago also acquires siDNAs during expression, we isolated and analyzed the nucleic acid fraction that co-purified with *Cb*Ago. Denaturing polyacrylamide gel electrophoresis revealed that *Cb*Ago co-purified with small nucleotides with a length of ~12-19 nucleotides (Figure 3A). These nucleic acids were susceptible to DNase I but not to RNase A treatment, indicating that *Cb*Ago acquires 12-19 nucleotide long siDNAs *in vivo*, which fits with its observed binding and cleavage activities *in vitro* (Figure 1 and 2).

**Figure 3.**
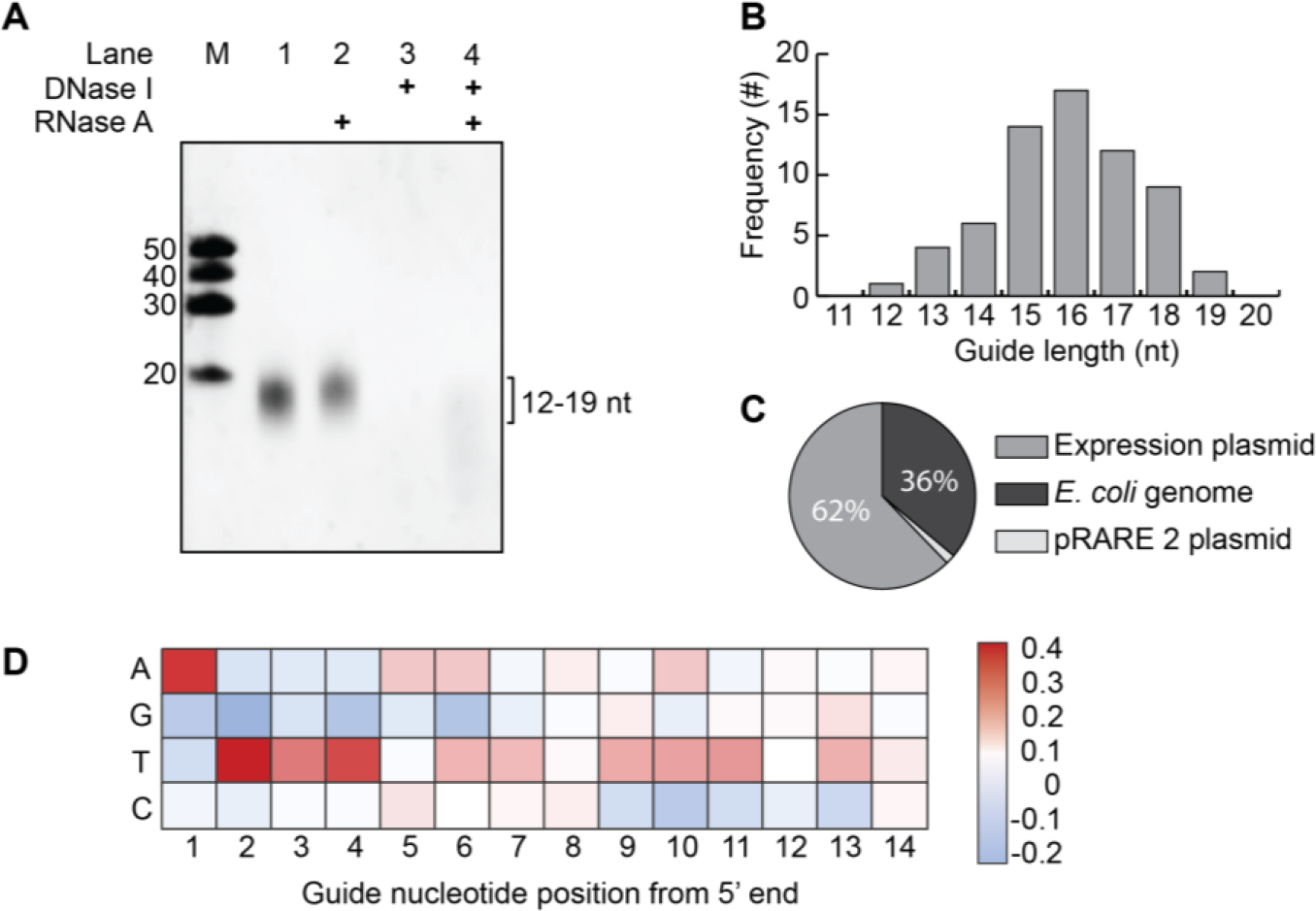
*Cb*Ago associates with small plasmid derived siDNA *in vivo*. (**A**) Nucleic acids that co-purified with *Cb*Ago were treated with either RNAse A, DNAse I or both, and were analyzed by denaturing polyacrylamide gel electrophoresis. (**B**) Histogram displaying the length of DNA co-purified with *Cb*Ago as determined by sequencing. (**C**) Sequenced nucleic acids that co-purified with *Cb*Ago are mostly complementary to the *Cb*Ago expression plasmid. (**D**) Heat map showing the base preference of the co-purified nucleic acids at each position. The red squares indicate bases that were more often found compared to a random distribution (25%); blue squares indicate bases that were less frequently found.

We cloned and sequenced the siDNAs that co-purified with *Cb*Ago to determine their exact length and sequence. The majority of the siDNAs had a length of 16 nucleotides and are complementary to the plasmid used for expression of *Cb*Ago (Figure 3B and 3C). Likewise the siRNAs and siDNAs that co-purify with respectively *Rhodobacter sphaeroides* (*Rs*Ago) and *Tt*Ago are also mostly complementary to their expression plasmids^13,16^. As both *Tt*Ago and *Rs*Ago have been demonstrated to interfere with plasmid DNA, this suggests that also *Cb*Ago might play a role in protecting its host against invading DNA. However, no significant reduction of plasmid content could be detected during or upon expression of *Cb*Ago in *E. coli* (Supplementary Figure S6). We also investigated whether *Cb*Ago co-purified with nucleic acids that were enriched for certain motifs. Sequence analysis revealed that most siDNAs co-purified with *Cb*Ago contain a deoxyadenosine at their 5’ ends (Figure 3D). In addition, we observed an enrichment of thymidine nucleotides in the three positions directly downstream of the siDNA 5’ end (nt 2-4) (Figure 3D).

### The sequence of the siDNA affects *Cb*Ago activity

To investigate if the 5’-terminal nucleotide of the siDNA affects the activity of *Cb*Ago, we performed cleavage assays. *Cb*Ago was loaded with siDNA guides with varied nucleotides at position 1 (g1N) and incubated with complementary target DNAs (Figure 4A). Surprisingly, the highest cleavage rates were observed when *Cb*Ago was loaded with siDNAs containing a 5’-T, followed by siDNAs containing 5’-A. *Cb*Ago bound 5’-G or 5’-C siDNAs displayed slightly lower initial cleavage rates. Also for other pAgos the g1N preference observed *in vivo* is not reflected in the *in vitro* activities; *Tt*Ago (which preferentially co-purifies with g1C siDNAs) as well as *Pf*Ago and *Mp*Ago (of which the *in vivo* g1N preferences are unknown) demonstrate no clear preference for a specific g1N during *in vitro* cleavage reactions^13,17,34^. Instead, the preference of *Tt*Ago for 5’-C siDNAs is determined by specific recognition of a guanosine nucleotide in the corresponding position (t1) in the target DNA^17^. Indeed, *Tt*Ago structures and models have revealed base-specific interactions with target strand guanine, while base-specific interactions with the 5’-terminal cytidine in the siDNA are less obvious^17^. Similarly, we observe no obvious base-specific interactions with the 5’-terminal cytidine in the structure of the *Cb*Ago complex (Supplementary Figure S7). When we investigated potential base-specific interactions with the base at the opposing target strand t1 position, we observed that the t1 thymine base is not placed in the t1 binding pocket as has been observed in *Tt*Ago, *Rs*Ago and hAGO2^17,39,40^. Instead, the thymine bases is flipped and stacks on Phe557 that also caps the siDNA-target DNA duplex (Supplementary Figure S7). At present, we are unable to rationalize the preferential co-purification of 5’-adenosine siDNAs with *Cb*Ago.

**Figure 4.**
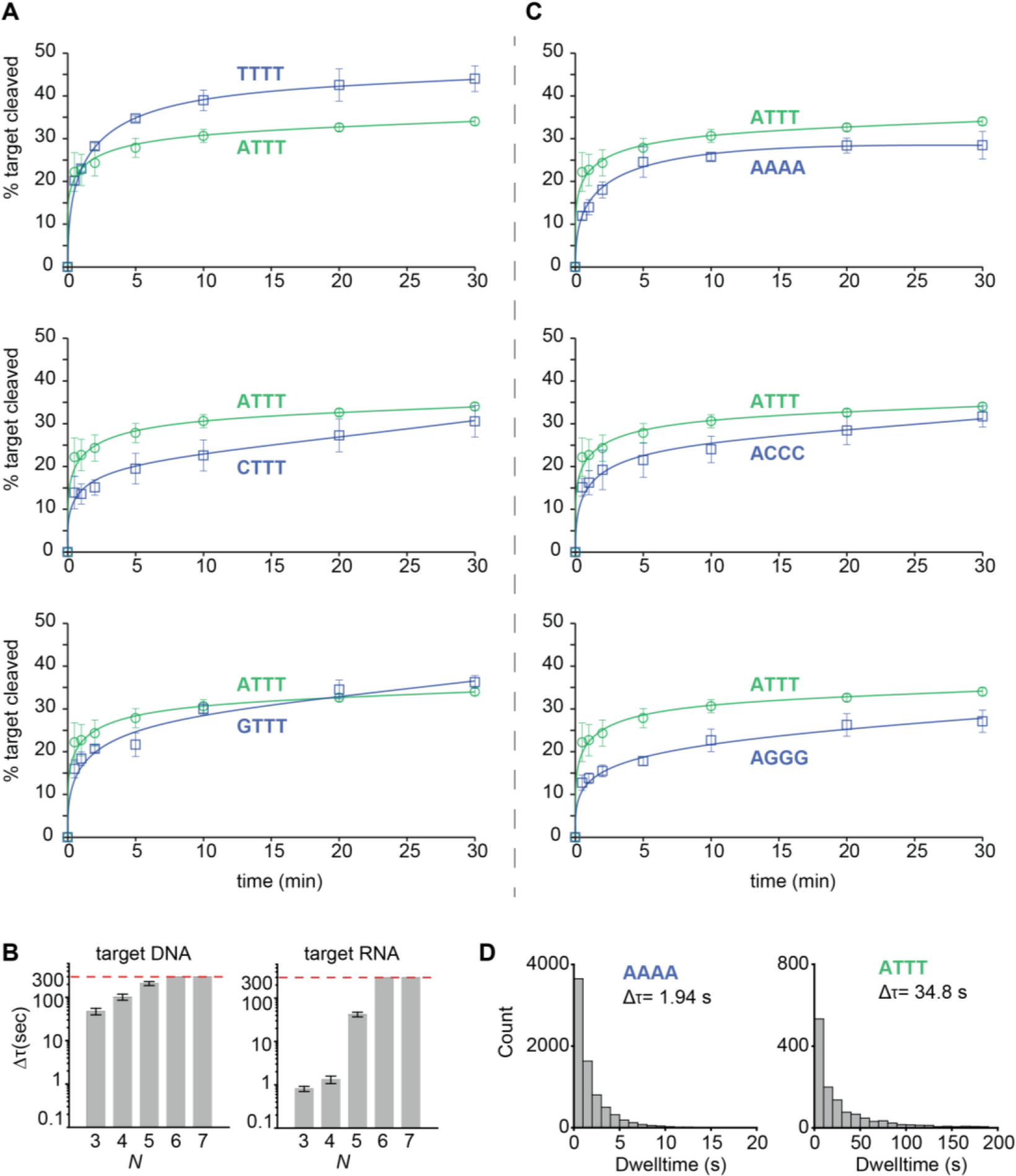
The siDNA sequence affects *Cb*Ago activity. (**A**) *Cb*Ago has no strong 5’-end nucleotide preference. *Cb*Ago was incubated with siDNA with varied 5’-end and incubated with complementary DNA targets. Cleavage products were analysed by denaturing polyacrylamide electrophoresis and quantified. Graphs display the amount of target DNA cleaved. Error bars indicate the standard variation of three independent experiments. (**B**) Histograms displaying dwell time of *Cb*AgoDM-siDNA complexes binding either DNA or RNA targets with a varied sequence complementarity (N = number of complementary nucleotides between the siDNA and the target, starting at nt2. Thus *N* 3= nt 2-4) The photobleaching limit is reached where the signal is deactivated (300s). (**C**) *Cb*Ago preferentially utilizes siDNAs with a TTT sub-seed segment. *Cb*Ago was incubated with siDNA with varied sub-seed segments (nt 2-4) and incubated with complementary DNA targets. Cleavage products were analysed by denaturing polyacrylamide electrophoresis and quantified. Graphs display the amount of target DNA cleaved. Error bars indicate the standard variation of three independent experiments. (**D**) Histograms displaying dwell time of *Cb*AgoDM in complex with a 5’-ATTT siDNA or 5’-AAAA siDNA binding to a target DNA. interactions that are on average ~18-fold longer than CbAgo in complex with siDNAs containing a 5’-AAAA motif.

In order to characterize the seed segment of *Cb*Ago, and to test whether the seed length changes depending on the nature of the guide and the target (*i.e.* DNA vs. RNA), we performed additional single-molecule binding assays. The length of seed was determined based on the minimal number of complementary nucleotide pairs between guide and target that were required to achieve a stable binding event. We first tested the sub-seed (nt 2-4), a 3-nt motif involved in initial target recognition in hAgo2^41,42^. When only the sub-seed segment of the siDNA is complementary to the DNA and RNA targets, *Cb*Ago-siDNA complexes bound to the DNA target with an average dwell time 58-fold longer compared to RNA target-binding (Figure 4B). When nt 2-7 of the guide were complementary to the target, the *Cb*Ago-siDNA complex stably bound to both to target DNA and RNA beyond our observation time of 300 s. This suggests *Cb*Ago prefers DNA targets above RNA targets and that the seed segment of the siDNAs bound by *Cb*Ago comprises nucleotides 2-7.

Next, we set out to investigate whether *Cb*Ago displays a preference for siDNAs with a TTT sub-seed (nt 2-4) *in vitro*, similar to the observed sequence preference for siDNAs that co-purified with *Cb*Ago *in vivo*. *Cb*Ago was incubated with siDNAs in which the sub-seed was varied and complementary target DNAs were added. In contrast to the 5’-base preference, the TTT sub-seed preference that we observed *in vivo* is also reflected *in vitro*: *Cb*Ago displays the highest target cleavage rates when programmed with TTT sub-seed siDNAs (Figure 4C). To confirm these findings, we performed single-molecule assays in which we compared the target binding properties of *Cb*Ago-siDNA complexes containing siDNAs with either a TTT or an AAA sub-seed segment. These assays demonstrate that the dwell time of *Cb*Ago loaded with a TTT sub-seed siDNA on a target was 18-fold longer compared to *Cb*Ago loaded with siDNA containing an AAA sub-seed (Figure 4D). Combined, these data indicate that *Cb*Ago displays a preference for siDNAs containing a TTT sub-seed segment.

### A pair of *Cb*Ago-siDNA complexes can cleave double stranded DNA

Thermophilic pAgos have successfully been used to generate double stranded DNA breaks in plasmid DNA^13,15^. As each pAgo-siDNA complex targets and cleaves a single strand of DNA only, two individual pAgo-siDNA complexes are required for dsDNA cleavage, each targeting another strand of the target dsDNA. Although all pAgos characterized so far appear to lack the ability to actively unwind or displace a dsDNA duplex substrate, it has been proposed that, at least *in vitro*, thermophilic pAgos rely on elevated temperatures (>65 °C) to facilitate local melting of the dsDNA targets to target each strand of the DNA individually. However, *Cb*Ago is derived from a mesophilic organism and we therefore hypothesize that it is able to mediate protection against invading DNA at moderate temperatures (37°C). To test if *Cb*Ago can indeed cleave dsDNA targets at 37°C, we incubated apo-*Cb*Ago and pre-assembled *Cb*Ago-siDNA complexes with a target plasmid. Previous studies showed that the ‘chopping’ activity of siDNA-free apo-*Tt*Ago and apo-*Mj*Ago can result in plasmid linearization or degradation, respectively^14,17^. We observed that apo-*Cb*Ago converted the plasmid substrate from a supercoiled to open-circular state, possibly by nicking one of the strands, but did not observe significant linearization or degradation of the plasmid DNA (Figure 5A). When the plasmid was targeted by *Cb*Ago loaded with a single siDNA, we also observed loss of supercoiling (Figure 5A). As this activity was not observed with nuclease-deficient *Cb*AgoDM, we conclude that apo-*Cb*Ago and *Cb*Ago-siDNA complexes are generate nicks in dsDNA plasmid targets with their DEDD catalytic site. When using two *Cb*Ago-siDNA complexes, each targeting one strand of the plasmid, we observed that a fraction of the target plasmid DNA becomes linearized (Figure 5A). This implies that *Cb*Ago-siDNA complex-mediated nicking of each of the target plasmid DNA strands resulted in the generation of a double stranded DNA break. Next, we investigated if the spacing between the two siDNAs affects the ability of *Cb*Ago to cleave the plasmid. The most efficient plasmid linearization was achieved when the siDNAs were orientated exactly or almost opposite to each other (Figure 5A).

**Figure 5.**
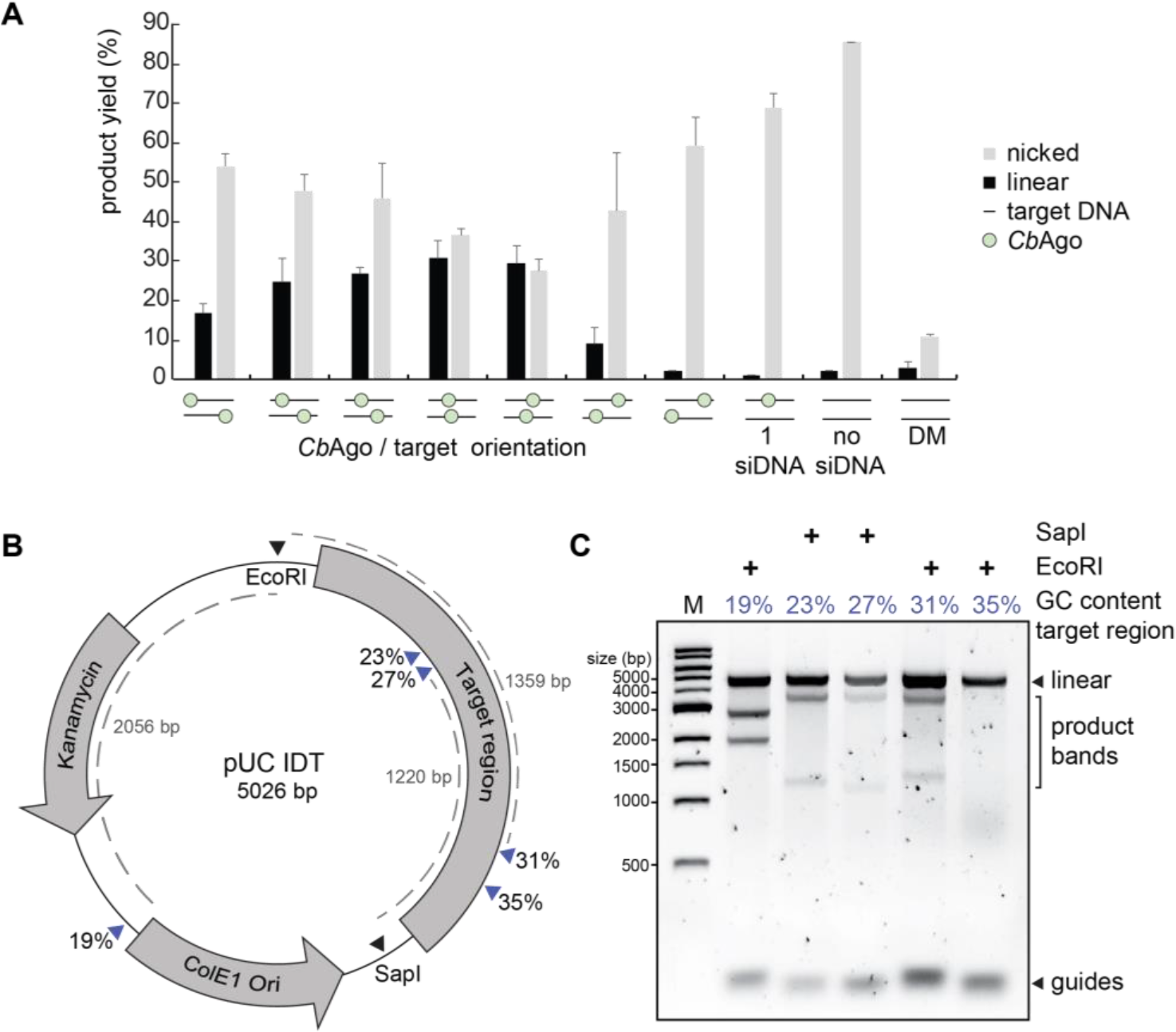
Double stranded plasmid DNA cleavage by *Cb*Ago. (**A**) Two *Cb*Ago-siDNA complexes can generate double stranded DNA breaks in plasmid DNA. CbAgo-siDNA complexes were pre-assembled and incubated with target plasmid DNA. Cleavage products were analysed by agarose gel electrophoresis and quantified. The spacing between both *Cb*Ago-siDNA target sites affects the linearization efficiency (nucleotide spacing between the predicted cleavage sites: +15 nt, +10 nt, +5 nt, 0 nt, −5 nt, −10 nt, −15 nt, a single siDNA, no siDNA). With 0 nt spacing, both *Cb*Ago-siDNA complexes are exactly on top of each other. (**B**) Schematic overview of the pUC IDT target plasmid. Blue arrows indicate target sites while percentages indicate the GC-content of the 100 bp segments in which these target sites are located. (**C**) Pre-assembled *Cb*Ago-siDNA complexes targeting various pUC IDT segments were incubated with pUC IDT. Cleavage products were incubated with EcoRI or SapI and were further analysed by agarose gel electrophoresis. The GC-content of the segments in which the target sites were located are indicated by the percentage (in blue),

Finally, we investigated whether the GC-content of the target DNA plays a role during DNA targeting by *Cb*Ago. For *Tt*Ago, it has been observed that AT-rich DNA is cleaved more efficiently than GC-rich DNA^17^. To test if such preference also exists for *Cb*Ago, we designed a target plasmid containing 16 gene fragments of 100 base pairs complementary to sequences from the human genome, with an increasing GC content (Figure 5B). *Cb*Ago-siDNA complexes were only able to generate dsDNA in gene fragments with a GC-content of 31% or lower (Figure 5C). This indicates that, at least *in vitro*, the GC-content is an important factor that determines target DNA cleavage by *Cb*Ago.

## Discussion

Several prokaryotic Argonaute proteins have been demonstrated to protect their host against invading nucleic acids, such as plasmid DNA^13,15,16^. Similar to *Tt*Ago and *Rs*Ago, *Cb*Ago co-purifies with guides which are preferentially acquired from the plasmid used for its heterologous expression in *E. coli*. In addition, *Cb*Ago mediates programmable DNA-guided DNA cleavage *in vitro*. This suggests that, similar to the phylogenetically related *Tt*Ago, also *Cb*Ago can interfere with plasmid DNA via DNA-guided DNA interference.

Sequencing of the nucleic acids that co-purified with *Cb*Ago revealed that *Cb*Ago preferentially associates with siDNAs with a 5’-ATTT-3’ sequence at their 5’ end. It was previously shown that the guide RNA utilized by eAgos can be divided into functional segments. These segments are (from 5’ to 3’) the anchor nucleotide (nt 1), the seed (nt 2-8) and sub-seed segments (nt 2-4), and the central (nt 9-12), 3’ supplementary (nt 13-16) and tail (nt 17-21) segments^41,43^. Extending this knowledge to the siDNAs that co-purified with *Cb*Ago, *Cb*Ago preferentially associates with siDNAs that have a 5’-terminal adenosine anchor (nt 1) and a T-rich sub-seed. In RNAi pathways, the preference for a specific 5’-terminal nucleotide is important for guide RNA loading into a subset of eAgos^44–46^ Similarly, several pAgos including *Rs*Ago, *Tt*Ago, and now *Cb*Ago also preferentially associate with specific 5’-terminal nucleotides *in vivo*^13,16^. However, for both *Cb*Ago and *Tt*Ago, there is no clear preference for siDNAs with that specific 5’-base during cleavage assays *in vitro*. Rather than having a functional importance, the preference of pAgos for a specific nucleotide at the siDNA 5’ end might be a consequence of siDNA generation and/or loading, as has been demonstrated for *Tt*Ago^17^. Several studies on human Ago2 have described the importance of the sub-seed segment (nt 2-4) in its RNA guides^41,42,47^. For hAgo2, a complete match between the guide RNA sub-seed segment and the target RNA triggers a conformational change that first exposes the remainder of the seed (nt 5-8), and eventually the rest of the guide. This facilitates progressive base paring between the guide RNA and the target^48^. However, a specific nucleotide preference in the sub-seed segment, as we have observed for *Cb*Ago, has not been described for any other Argonaute protein. The preference for the T-rich sub-seed is not only observed in the *in vivo* acquired siDNAs, but also plays a clear role during target binding and cleavage assays *in vitro.* This may reflect a structural preference for these thymidines in the cleft of the PIWI domain. We have not been able to obtain diffracting crystals of *Cb*Ago in complex with siDNAs that have a 5’-ATTT-3’ sequence at the 5’-end. Future research will thus be necessary to determine the structural basis the apparent preference for these nucleotides at these positions. We hypothesize that this bias might reflect the mesophilic nature of *Cb*Ago, which might have better access to AT-rich dsDNA fragments, both for siDNA acquisition and for target cleavage.

Several DNA-targeting pAgos have been repurposed for a range of molecular applications among which a cloning, recombineering and nucleic acid-detection method^22,23,49,50^. Additionally, the potential repurposing of pAgos for genome editing applications has previously been discussed^27^. However, all characterized DNA-cleaving pAgos to date originate from thermophilic prokaryotes and are solely active at elevated temperatures, which limits the potential repurposing of pAgos for applications that require moderate temperatures, such as genome editing. The biochemical characterization of *Cb*Ago reported herein is the first example of a pAgo that catalyzes siDNA-guided dsDNA cleavage at 37°C, indicating that the pool of mesophilic pAgos contains candidates that – in theory – can be utilized for potential applications that require moderate temperatures, such as genome editing. If *Cb*Ago or other mesophilic pAgos could be harnessed for genome editing, they will have certain advantages over the currently well-established genome editing tools CRISPR-Cas9 and CRISPR-Cas12a; While CRISPR-based genome editing tools can be programmed with a guide RNA to target DNA sequences of choice, target DNA cleavage additionally requires the presence of a protospacer adjacent motif (PAM) next to the targeted sequence (5’-NGG-3’ for Cas9 and 5’-TTTV-3’ for Cas12a)^51^. This limits the possible target sites of Cas9 and Cas12a. In contrast, pAgos do not require a PAM for DNA targeting, which would make them much more versatile tools compared to CRISPR-associated nucleases. However, PAM binding by Cas9 and Cas12a also promotes unwinding of dsDNA targets^52–54^ which subsequently facilitates strand displacement by the RNA guide, and eventually R-Loop formation. The absence of such mechanism in pAgos might explain their limited nuclease activity on dsDNA targets.

Here, we have demonstrated that *Cb*Ago does not strictly rely on other proteins when targeting AT-rich dsDNA sequences *in vitro*. As such, this study provides a foundation for future efforts to improve double stranded DNA target accessibility of pAgos and to facilitate the further development of pAgo-based applications at moderate temperatures.

## Acknowledgements

We are grateful to Meitian Wang, Vincent Olieric, and Takashi Tomizaki at the Swiss Light Source (Paul Scherrer Institute, Villigen, Switzerland) for assistance with X-ray diffraction measurements. This work was supported by grants from the Netherlands Organization of Scientific Research (NWO; ECHO grant 711013002 and NWO-TOP grant 714.015.001) to J.v.d.O. A Swiss National Science Foundation (SNSF) Project Grant to M.J. (SNSF 31003A_149393) and by long-term postdoctoral fellowships from the European Molecular Biology Organization (EMBO) to D.C.S (ALTF 179-2015 and aALTF 509-2017). M.J. is International Research Scholar of the Howard Hughes Medical Institute and Vallee Scholar of the Bert L & N Kuggie Vallee Foundation. C.J. was supported by Vidi (864.14.002) of the Netherlands Organization for Scientific research.

## Author contributions

J.W.H. and J.v.d.O. conceived the project and designed the biochemical experiments, which were performed by J.H and J.K. Single-molecule experiments were designed by S.C., T.J.C and C.J. and performed by S.C and T.J.C. X-ray crystallographic analysis was designed and performed by D.C.S. under the supervision of M.J.. J.W.H., D.C.S., C.H., M.J., C.J. and J.v.d.O. wrote the manuscript. All authors read and approved the manuscript.

